# The coloNISation: spatio-temporal metabarcoding surveys in ports reveal homogenised communities with high genetic diversity and connectivity of non-indigenous species

**DOI:** 10.1101/2025.11.18.688838

**Authors:** J. Zarcero, A. Antich, M. Fernández-Tejedor, C. Palacin, O.S. Wangensteen, M. Rius, X. Turon

## Abstract

Large commercial ports facilitate the introduction of non-indigenous species (NIS), while smaller harbours and marinas contribute to their regional spread. Harbour networks are thus important drivers of introductions. Despite extensive research effort on NIS in recent years, no study has yet assessed genetic connectivity among harbours considering whole-community composition. Here, we analysed spatio-temporal patterns of metazoan communities over one year in four medium-size harbours along the NW Mediterranean coast sampled by deploying standardised biological collectors. Using cytochrome c oxidase subunit I (COI) metabarcoding, we identified 1,770 metazoan molecular operational taxonomic units (MOTUs), of which 82 were classified as NIS based on a custom database of Mediterranean NIS. Despite their lower species count compared to natives, NIS accounted for 34–70% of reads in harbours. The southernmost harbour had the highest NIS number of reads, likely due to its proximity to aquaculture facilities. While we observed some variation in the spatial structure of metazoan communities across harbours, NIS showed consistently low differentiation values, sharing significantly more MOTUs among sites. Seasonal patterns influenced both NIS and the rest of the community. Haplotype diversity was significantly higher in NIS, which also exhibited lower genetic differentiation across harbours compared to native species, indicating NIS spread via local boating and likely recurrent introductions. These findings highlight distinct dynamics between NIS and native species in artificial environments, emphasising the importance of continued monitoring in harbour networks to manage coastal NIS proliferation.

**HIGHLIGHTS:** COI metabarcoding of standardised collectors detected over 1,700 MOTUs of marine metazoans in ports over a year.

Less than 4% were NIS MOTUs, but they comprised 34-71 % of the reads.

NIS were more homogeneously distributed among ports than other MOTUs.

NIS showed higher genetic variability but lower genetic differentiation than native species.

Different dynamics underpin NIS and native assemblages in port communities.

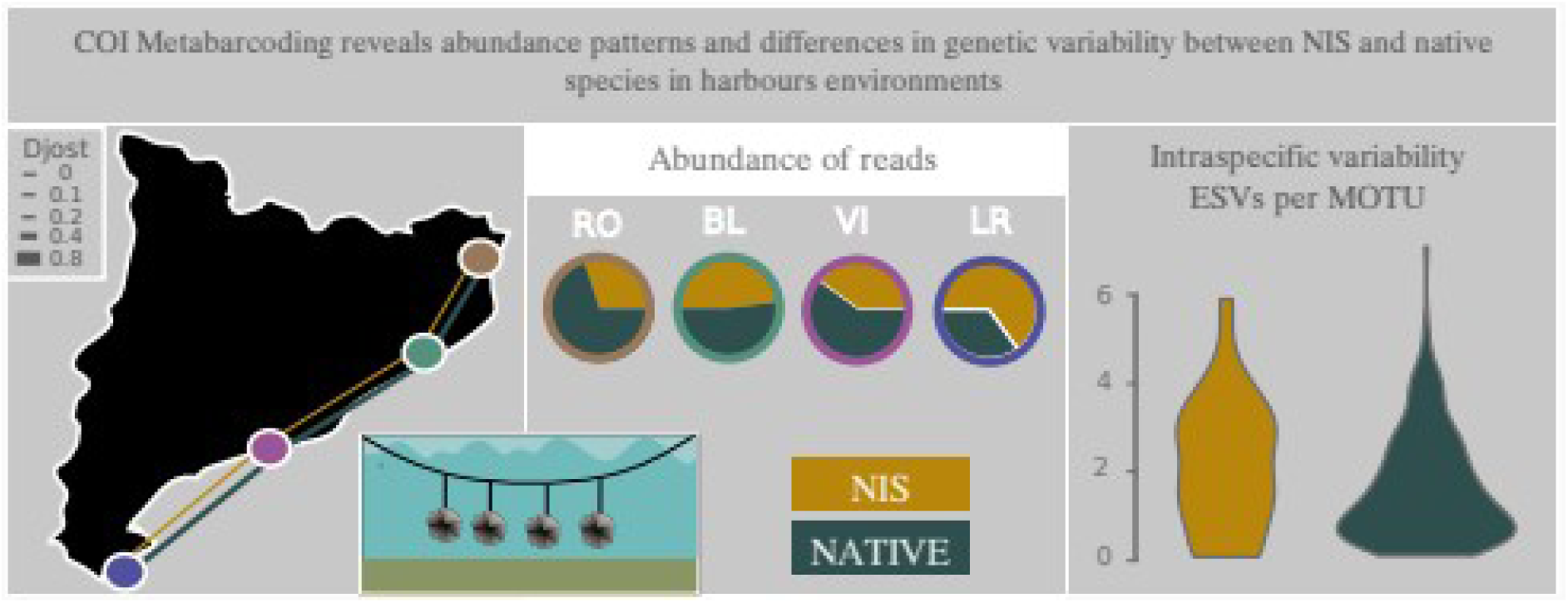

## 1. INTRODUCCION

Population connectivity is a key process shaping community structure (Cowen et al. 2009; Treml et al. 2015). Although most studies of population connectivity focus on natural habitats, increasing attention has been given to connectivity among highly impacted, anthropised ecosystems (Preston et al. 2025). These studies often focus on a single or a small group of species (Melià et al. 2016) and thus little is known about population connectivity among these ecosystems considering whole community composition.

Ports and marinas are emerging as ubiquitous elements of coastal seascapes, being part of the so-called coastal urban sprawl (Bishop et al., 2017). The continuous expansion of these artificial infrastructures profoundly impacts natural littoral communities and raises significant socio-ecological concerns (Floerl et al., 2021). Differences in biotic assemblages between natural and artificial habitats have been widely documented (Bonnici et al., 2018; Madon et al., 2023; Mayer-Pinto et al., 2018; Pagnier et al., 2024). However, ports offer a unique setting for studying evolutionary responses to both novel population connectivity networks and changing environmental conditions (Touchard et al., 2023, Lilli et al., 2025). Despite a recent focus on research in these ecosystems, there is a lack of studies focussing on spatio-temporal assessments of community structure in port environments (Bishop et al., 2017).

Ports also serve as hotspots for the establishment and spread of non-indigenous species (NIS), which are a growing concern for a wide array of ecosystems (e.g., Macé et al., 2024; Madon et al., 2023; Mineur et al., 2012; Miralles et al., 2021). Port communities are typically composed of species tolerant to changing biotic conditions, pollution, and other harbour-specific stressors. Propagule pressure as a result of maritime traffic is a major driver of NIS introductions (Andrés et al., 2023; Lacoursière-Roussel et al., 2016), contributing to the richness of NIS in the Mediterranean Sea (Spagnolo et al., 2019; Ulman et al., 2019a, 2019b). As such, ports are central to NIS dynamics and warrant focused studies (Airoldi et al., 2015; Bishop et al., 2015; Gewing and Shenkar, 2017; Tempesti et al., 2020). While previous genetic studies on NIS in artificial environments have typically adopted a species-by-species approach (e.g., Hudson et al., 2016; Ordóñez et al., 2013; Rius et al., 2012; Dupont et al., 2010; Pineda et al., 2016; Zardus & Hadfield, 2005), community-level comparisons that include both native and NIS have the potential to shed light on the metacommunity-level population connectivity of harbour ecosystems.

Although taxonomy-based field surveys greatly aid NIS detection in ports (e.g., Ammon et al., 2018; Arenas et al., 2006; Cohen et al., 2005), species identifications can be time-consuming, often require specialised expertise, and can be particularly complex in taxonomically challenging groups such as ascidians, bryozoans, and cnidarians (Viard et al., 2019; Duarte et al., 2021; Pagenkopp Lohan et al., 2016; Martins et al., 2021). Consequently, molecular methods, especially metabarcoding, are increasingly used for port community monitoring (Alexander et al., 2023; Holman et al., 2019; Lavrador et al., 2024). These tools generate high-throughput biodiversity data that can reveal whole-community composition, detect NIS, and monitor temporal change (Rey et al., 2020). Nonetheless, challenges remain for metabarcoding studies, such as methods’ standardisation and the choice of markers and sampling substrates (Alexander et al., 2023; Deiner et al., 2018; Lavrador et al., 2024). Thus, further development of reference databases and bioinformatic pipelines are crucial for enhancing the reliability of metabarcoding in NIS-focused harbour studies (Couton et al., 2022). Adequate geographic and temporal coverage is also critical for assessing connectivity and seasonal trends (Hoedt et al., 2001; Rey et al., 2020; Lavrador et al., 2024).

Genetic and genomic studies at intraspecific level, typically targeting one or a few NIS (e.g., Galià-Camps et al., 2023; Hudson et al., 2022; Ordóñez et al., 2013; Turon et al., 2003), provide insights into invasion processes such as the so-called genetic paradox: invasive species often succeed despite reduced genetic diversity resulting from bottlenecks during introduction (Estoup et al., 2016; Golani et al., 2007). Multiple introductions may counteract this, enhancing genetic richness and promoting anthropogenic homogenisation (Hudson et al., 2022; Roman and Darling, 2007). Understanding NIS genetic diversity can therefore shed light on their invasion dynamics and the impacts on native species (Stepien et al., 2002). Metabarcoding data also allow for metaphylogeographic analyses that allow extracting intraspecific genetic signals such as haplotype composition (Turon et al., 2020), and computing genetic differentiation measures among populations from denoised sequence variants (Antich et al., 2021, 2022, Riley et al., 2025). This enables comparisons of genetic richness and connectivity between native and NIS’ populations, helping to explain differences in fitness, adaptability, and dispersal success (Pauls et al., 2013; Wade et al., 2017).

In this study, we analysed metazoan communities across four similarly sized ports with fishing and commercial activities. We specifically avoided large commercial ports to concentrate on post-border processes - i.e., secondary spread via local boating and fishing (Forrest et al., 2009; Darbyson, 2009; Davidson et al., 2010; Outinen et al., 2021). By examining community structure and comparing native versus NIS dynamics, we explored both interspecific and intraspecific patterns (in line with Turon et al., 2020), with special attention to unravelling patterns of genetic diversity and population connectivity. Our general expectations are that ports act as networks facilitating the connectivity of NIS (Airoldi et al 2015, López-Legentil et al 2015) and that recreational boating between ports leads to recurrent introduction of NIS. We specifically tested the following working hypotheses: 1) NIS will have higher homogeneity (i.e., lower beta-diversity) between ports than the overall community; 2) NIS will exhibit higher population connectivity and less genetic differentiation than native species and 3) NIS will display greater levels of genetic diversity within port communities compared to native species. Our approach will allow assessing how dispersal dynamics and propagule pressure influence the invasion processes and the resilience of native species.

## 2. MATERIAL & METHODS

### 2.1. Sampling and sample processing

We focused on the Catalan coastline (NW Mediterranean), a densely urbanised region with a high concentration of ports (López-Legentil et al., 2015). We used a recently designed sampling device called POMPOMs (Zarcero et al., 2024), which consists of a 100 × 25 cm strip of polyamide mesh with hexagonal openings of approximately 1 mm, folded in a zig-zag pattern and fastened with cable ties to form a circular shape (approximately 25 cm in diameter). POMPOMs are capable of capturing both early life-history stages and particulate organic matter, thereby providing a comprehensive snapshot of the biodiversity present.

POMPOMs were placed underwater (ca. 1 m deep) hanging from dedicated ropes in the ports. Four replicate collectors were deployed in four ports on the Catalan coast (northwestern Mediterranean): Roses (42.254495 °N, 3.180719 °E), Blanes (41.674425 °N, 2.799687 °E), Vilanova i La Geltrú (41.214873 °N, 1.736144 °E) and La Ràpita (40.618726 °N, 0.598968 °E) (see details in Fig. S1). We will hereafter denote these locations as RO (Roses), BL (Blanes), VI (Vilanova I la Geltrú) and LR (La Ràpita). All these ports are medium-sized and sustain fishing and recreational boating activity. They comprise altogether between 1,500 to 3,000 lineal metres of docks. The distance by sea between the two most distant ones (RO and LR) is 323 Km.

The POMPOMs were deployed from February 2019 to April 2020. They were replaced monthly during warm months and bimonthly during cold months, resulting in a total of 10 temporal sampling points. Three collectors per month and site were used for metabarcoding studies, while the fourth was retained as a backup. In total, 120 samples were obtained for this study. After collection, the POMPOMs were unfolded and all biofouling was carefully removed using sterilised 10 cm nylon brushes and recovered in a stainless-steel sieve of 64 µm.

We conducted seawater temperature readings at each site every hour over the study period using HOBO^(R)^ Pendant data loggers (resolution 0.53°C), which were placed right next to the POMPOMs (see details in Fig. S2).

### 2.2. DNA extraction & sequencing

All procedures were performed in a sterilised laminar flow cabinet, with UV light activated between each sample processing. We followed the DNA extraction, library preparation, and sequencing protocols detailed in Antich et al. (2023). In short, DNA was extracted from 5 g of homogenised sample material using the DNeasy PowerMax Soil Kit (Qiagen). A fragment of ca. 313 bp of the cytochrome c oxidase subunit I (COI) gene was amplified using as generalist primers for eukaryotes the Leray-XT primer set (Wangensteen et al., 2018), which includes the forward primer jgHCO2198: 5’-TAIACYTCIGGRTGICCRAARAAYCA-3’ (Geller et al., 2013) and the reverse primer mlCOIintF-XT: 5’-GGWACWRGWTGRACWITITAYCCYCC-3’. Both primers were tagged with an 8-base sequence at the 5’ end, with distinct tags for each sample differing by at least three bases. The same tag was used for both forward and reverse primers of each sample to facilitate the elimination of inter-sample chimeras. In order to enhance sequence diversity and facilitate Illumina base calling, a variable number of degenerate bases (N), from two to four, were added before the tags on both primers. The PCR procedure consisted of a first denaturation step for 10 min at 95 °C, followed by 45 cycles of denaturation at 94 °C for 60 s, hybridisation at 45 °C for 60 s and elongation at 72 °C for 60 s, ending with a final elongation step of 5 min. The amplification products were then purified and concentrated using the MinElute PCR Purification Kit (Qiagen). The success of the amplification was verified using gel electrophoresis. Negative samples (n=12) were run by processing and extracting samples of sand charred in a muffle furnace. Fifteen PCR blanks, containing no DNA template, were also included. Libraries were prepared using the BIOO NEXTFLEX PCR-Free DNA-Seq Kit (Perkin-Elmer) and sequenced on a partial Illumina NovaSeq lane with 2 x 250 bp paired-end sequencing at Novogene Company.

### 2.3. Bioinformatic analyses

We used a custom pipeline based on the Obitools3 software (Boyer et al., 2016) and using both Bash and R 4.0.2 scripts. Briefly, Illuminapairedend was used to align paired-end reads, retaining only those reads with an alignment quality score greater than 40. Reads were then demultiplexed using ngsfilter, discarding any reads with unmatched primer tags at their ends. The obigrep and obiuniq functions were used to perform a length filter (retaining only sequences between 310 and 319 bp) and combine identical sequences. Singleton sequences (with just one read) were deleted at this step. The Uchime de novo algorithm of VSEARCH v2.7.1 was then used to remove chimeric amplicons (Rognes et al., 2016).

Sequences were denoised with the DnoisE program (Antich et al., 2022), which is a modification of the Unoise algorithm (Edgar, 2016) that incorporates the natural variability in the three codon positions of coding genes. Denoising was performed within samples with an alpha parameter of 4 and an auto-computed entropy correction to generate Exact Sequence Variants (ESVs) (Antich et al., 2021). The following filters were then applied to the ESV dataset: We first deleted any ESV for which reads in blanks or negative controls represented more than 10% of total reads for that ESV in all samples, as these are suspected to correspond to contaminations. Second, for each sample, a dual abundance filtering was established, setting to zero the reads of (i) ESVs that represented less than 0.005% of the sample total reads, and (ii) ESVs with less than 5 reads after the previous step. ESVs were then clustered into molecular operational taxonomic units (MOTUs) with SWARM v3.1.3 using d = 13, following Antich et al., (2021). SWARM is a variable-threshold and fast algorithm that connects all reads with a distance less than d in a first step and then breaks down the resulting clusters using a topological criterion based on the internal abundance structures of the clusters (Mahé et al., 2014, 2022). The most abundant ESV in the MOTU was used as the representative sequence. The information of which MOTU each ESV was assigned to was also extracted from the SWARM output and added to the ESV dataset.

Taxonomic assignment of the representative sequences of the MOTUs was performed using the mkLTG software (Meglécz, 2024) with the COInr database (Méglecz, 2023) . The default parameters of the mkLTG procedure were modified (the updated configuration is available at https://github.com/jesuszarcero/mkLTG_params_modified). To avoid overclassification (Mugnai et al., 2023), we eliminated Insecta and Arachnida (but keeping Acari) from the reference database using the mkCOInr program (Meglécz, 2023). Only MOTUs assigned to Metazoa were kept for downstream analyses. The final dataset refinement consisted of applying to the MOTUs dataset the LULU post-clustering correction procedure (Frøslev et al., 2017). This procedure combines similarity and co-occurrence metrics to detect erroneous MOTUs and pools the reads with the correct ones. The ESV dataset was updated accordingly. We used a modification of the original LULU function (https://github.com/jesuszarcero/LULU_corrected) to correct a bug previously detected (https://github.com/tobiasgf/lulu/issues/8). A final checking of erroneous sequences, including nuclear mitochondrial inserts (numts) was performed following Turon et al., (2020) and Zarcero et al., (2024): all ESVs whose representative sequences had codon stops were deleted, after checking the 12 metazoan mitochondrial genetic codes from the Biostrings R package (Pagès, 2017). We also checked the five amino acids conserved across metazoans in the studied fragment (Pentisaari et al., 2016), and sequences with changes in these positions were deemed as incorrect and the corresponding ESV deleted. If all ESVs of a MOTU were eliminated, the latter was also deleted from the final MOTU table.

NIS detection was performed using the NCBI-BLAST algorithm (Korf et al., 2003) and the custom NISdb v3 database (Zarcero et al., 2024) (https://github.com/jesuszarcero/NISdb), which contains curated sequences of NIS found in the Mediterranean Sea. The NIS database was compiled based on the available literature on NIS in the Mediterranean and is regularly updated. Subsequently, we searched for available COI-5P sequences of these species in the Barcode of Life Data System (BOLD, https://www.boldsystems.org/, accessed in October 2023). Whenever NIS sequences were not found in BOLD, we referred to the NCBI database and verified species identification using the taxonomic literature. For each NIS, the obtained sequences were collapsed into unique haplotypes. To ensure data accuracy, a thorough manual curation was conducted, involving comprehensive BLAST searches and the removal of potentially erroneous sequences whenever the BLAST results indicated a different species or conflicting assignments. BLAST results were filtered, retaining assignments with an identity equal to or greater than 97% and a sequence coverage equal to or greater than 70%. Assignments against this database were compared with the corresponding MOTU assignments obtained with the mkLTG procedure to check the accuracy of NIS detection with the general database.

### 2.4. Datasets

Downstream analyses were conducted using three different datasets, depending on the objective of each analysis. We first identified MOTUs assigned to non-indigenous species (NIS) following the procedure described above, and the remainder of the community (hereafter the COMM dataset). This dataset includes many MOTUs that could not be identified to species level; therefore, their exact status as native or NIS could not be determined, albeit we assume that most of them were native, as genetic databases for NIS are more complete (Jerde et al., 2021). To enable more focused comparisons with the NIS dataset, we also selected those COMM MOTUs identified to species level and confidently assigned as native (hereafter NAT). The NAT list was manually curated to exclude problematic MOTUs, such as species complexes, dubious database matches, or cryptogenic species.

### 2.5. Statistical analyses

Unless otherwise stated, analyses were conducted with the ‘vegan’ R package (Oksanen et al., 2019), and plots were created with the ‘ggplot2’ R package (Wickham, 2011). Rarefaction curves and species accumulation curves were generated with the functions rarecurve and specaccum, respectively. MOTU richness and Shannon diversity values were computed after rarefaction to the minimum number of reads in any sample using the function rarefy. The resulting metrics were compared across categories with ANOVAs and post hoc Tukey tests.

To assess the taxonomic composition of the samples, MOTUs were grouped into major metazoan phyla, and bar charts were generated to display the average composition per locality in terms of proportion of reads and proportion of MOTUs for both the COMM and the NIS datasets. The proportion of reads assigned to each phylum was also broken down by month at each locality to ascertain temporal trends. Additionally, a heatmap was constructed to visualise the prevalence of each NIS across ports. Upset plots were used to visualise patterns of MOTU sharing across localities for the COMM and NIS datasets, and these patterns were summarised by computing a match index (Ahrens et al., 2016) with following equation:

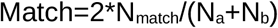

Where N_match_ is the number of shared MOTUs between sites a and b, and N_a_ and N_b_ are the total number of MOTUs in each of them, respectively.

For β-diversity analysis, we used the relative read abundance of each MOTU in each sample without rarefaction and computed the Bray-Curtis dissimilarity index (BC). These values were then used to generate reduced-space representations of the samples in a non-metric multidimensional scaling (NMDS) configuration using the metaMDS function of vegan, separately for COMM and NIS MOTUs.

Permutational analyses of variance (PERMANOVA) were conducted for the factors site and month on the BC matrix using the PERMANOVA module of the statistical package Primer v6 (Anderson and Walsh, 2013). Pairwise tests for significant factors were performed. A total of 999 permutations were used to generate the statistics of these analyses. We also performed dispersion tests (permdisp procedure with 999 permutations) to assess differences in multivariate dispersion of the groups of samples. Finally, we analysed temporal trends of community change by averaging Bray-Curtis dissimilarities among consecutive sampling times, separately for the COMM and NIS datasets.

### 2.6. Metaphylogeographic analyses

We also performed analyses of genetic diversity within MOTUs (our proxy for species-level diversity) using the ESVs as a proxy for their haplotype composition (Turon et al., 2020). For this assessment, we restricted our analyses to comparisons of the NIS and NAT datasets to ensure a correct assignment to these categories, as the COMM dataset had many MOTUs unassigned at species level. We first obtained the values of ESV richness in NIS and NAT MOTUs and visualised them in violin plots. Given their highly skewed nature, they were compared via a non-parametric rank test (Mann-Whitney’s U).

We then computed genetic differentiation measures among pairs of sampling localities using the D statistic (Jost, 2008) with the function *pairwise_D* of the R package ‘mmod’ (Winter, 2012). We selected, for each pair of localities, those shared MOTUs with at least 2 ESVs in the localities in order to ensure a minimal level of genetic variation. Read frequency data do not provide a direct measure of abundance (Turon et al., 2020), thus an indirect estimate was used, following Antich et al., (2023) and Azarian et al. (2020). We considered as an abundance approximation the frequency of occurrence of each ESV, that is, the number of samples where each ESV has been detected in each locality. Occasional negative D values were transformed to zeros. These analyses were performed separately for MOTUs assigned to NIS and NAT, and Jost’s D values were compared among pairs of localities with paired-sample t-tests.

## 3. RESULTS

### 3.1. Global diversity and datasets

All sequences generated were uploaded to the NCBI SRA archive (Bioproject PRJNA1179718). After pairing, demultiplexing, quality and length filtering, and chimera removal, we obtained 283,820,787 reads from 3,013,217 unique COI sequences. The denoising procedure resulted in 41,792 ESVs, which were grouped into 9,006 MOTUs. After all filtering steps, 4,260 MOTUs and 24,782 ESVs were kept. From these, we retained all those assigned to marine metazoans. Ten samples were left with less than 9,500 reads and were discarded. Our final database thus consisted of 110 samples with a total of 1,774 MOTUs, 11,193 ESVs, and 123,883,837 reads. The final MOTU table, with taxonomic identification and assignment to the different datasets, is presented in Table S1. The final ESV table, with indication of the MOTU to which each ESV belongs, is given as Table S2. The rarefaction curves (Fig. S3A) showed that a plateau was reached in the number of MOTUs in the samples, indicating adequate sequencing depth. Conversely, the MOTU accumulation curves (Fig. S3B) did not reach a plateau, with more MOTUs being added as more samples were included.

We detected 82 MOTUs belonging to NIS using BLAST with the NISdb, comprising 68,735,542 reads. These reads were distributed across 57 nominal species (Table S1). When we checked the taxonomy obtained with NISdb with the one obtained from the COInr database with mkLTG, they were coincident in 73 MOTUs, while for 7 MOTUs the mkLTG procedure could assign them only at genus or higher levels, for 2 MOTUs the mkLTG assignment was to a different species, and for another three it was to the same species but with less than 97% identity. The COMM dataset comprised the remaining 1,694 MOTUs and 55,148,295 reads. Of these, 237 MOTUs assigned at the species level were identified as native, with 24,301,437 reads and belonging to 212 nominal species (Table S1).

### 3.2. Taxonomic composition

Bar charts were prepared to show the overall taxonomic composition separately for each locality in terms of metazoan phyla composition for the COMM and NIS datasets (Fig. 1). The proportions were calculated based on the averaged relative abundances of the reads per locality. Phyla representing less than 5% of the reads were grouped under "Others”. The "Unidentified" category corresponded to metazoan reads that could not be assigned to phylum level. The group with the highest relative abundance in the COMM dataset were the arthropods, with cnidarians second. These phyla showed contrasting relative abundance trends, increasing from North to South for the former and decreasing for the latter. The third most abundant group were chordates (mostly ascidians), followed by annelids, molluscs, bryozoans, and “Others”. The “Unidentified” category had between 2% and 60% of the reads across all samples. When considering the NIS dataset, arthropods, annelids, cnidarians and chordates were, in this order, the dominant phyla in terms of relative read abundance, with marked differences across ports. For instance, BL had a higher abundance of annelids and chordates, while arthropods were the dominant group in the other ports, particularly RO and LR (>70% of NIS reads).

**Fig. 1.**
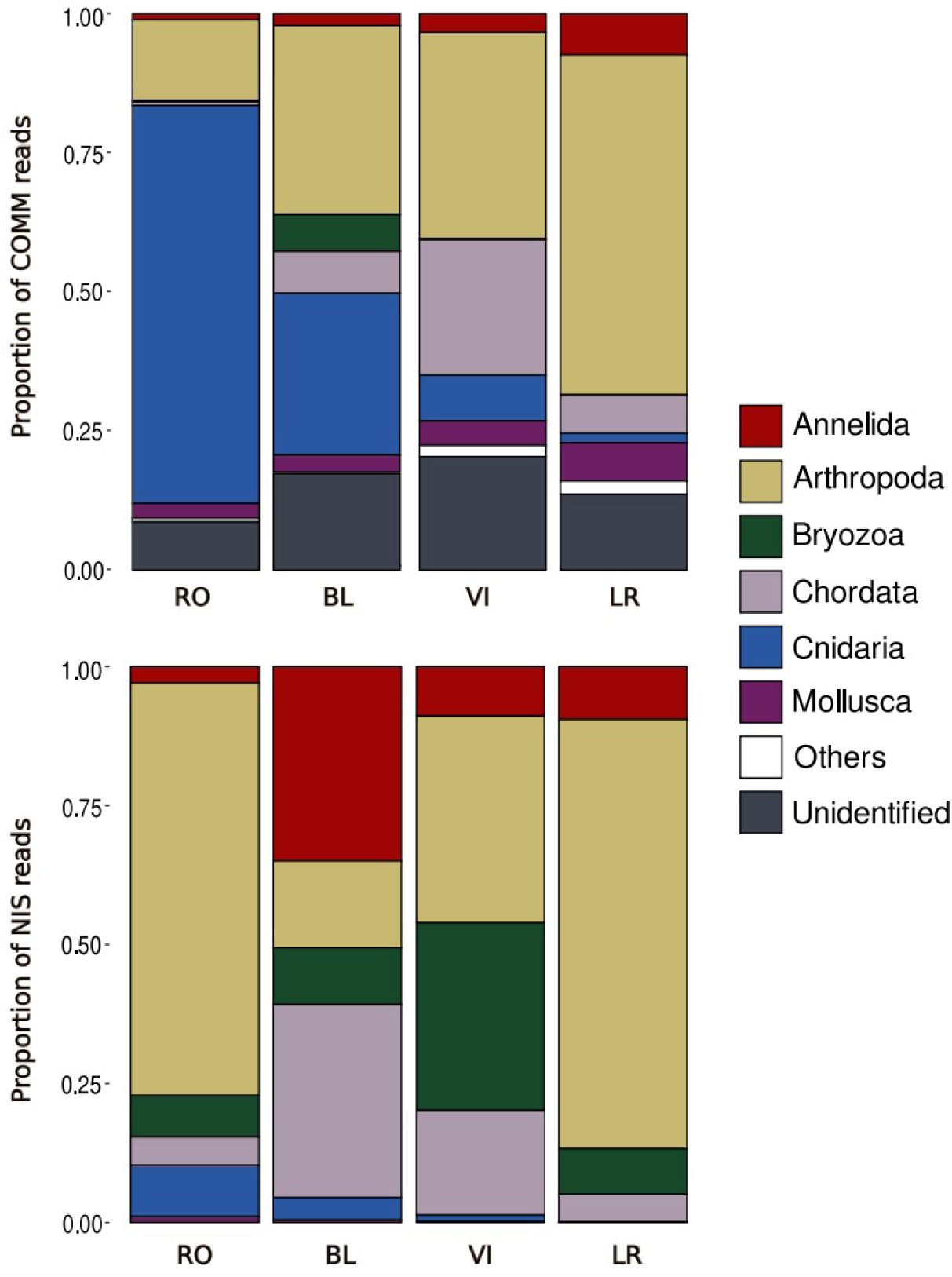
Barplots of the relative proportion of COMM (A) and NIS (B) reads of the different phyla for each area (RO: Roses; BL: Blanes ; VI: Vilanova i La Geltrú; LR: La Ràpita).

The taxonomic composition of the samples is presented separately for each sampling time for the COMM (Fig. S4) and NIS (Fig. S5) datasets. For the COMM dataset, in north ports (RO and BL), there was a trend of dominance of arthropods in spring-beginning summer, with higher relative abundance of cnidarians in late summer to winter. In turn, southern ports (VI and LR) showed that arthropods were in general the best represented group all year round (with the exception of July 2019), and cnidarians were not abundant, except in May 2019 in VI. The time course of relative abundances in the NIS dataset did not show a common pattern, with arthropods dominating most of the year in the northernmost (RO) and southernmost (LR) sites, and a strong presence of annelids in summer-fall in BL, and of bryozoans in fall-winter in VI. Chordates (mostly ascidians) were particularly abundant in BL and VI, but did not show a clear temporal pattern.

### 3.3. ɑ-diversity

The mean richness and Shannon diversity values after rarefaction (at 9,751 reads, the minimum number of reads of any sample) are shown in Fig. 2. For both metrics, BL had the highest values in the COMM dataset and the lowest values in the NIS dataset. For the COMM dataset, the mean diversity values were 2.76 ± 0.42 overall and the richness mean was 98.4 ± 40.1 MOTUs. For the NIS dataset, the mean diversity value for the rarefied data was 1.84 ± 0.35, and the mean richness value was 19.32 ± 4.70.

**Fig. 2.**
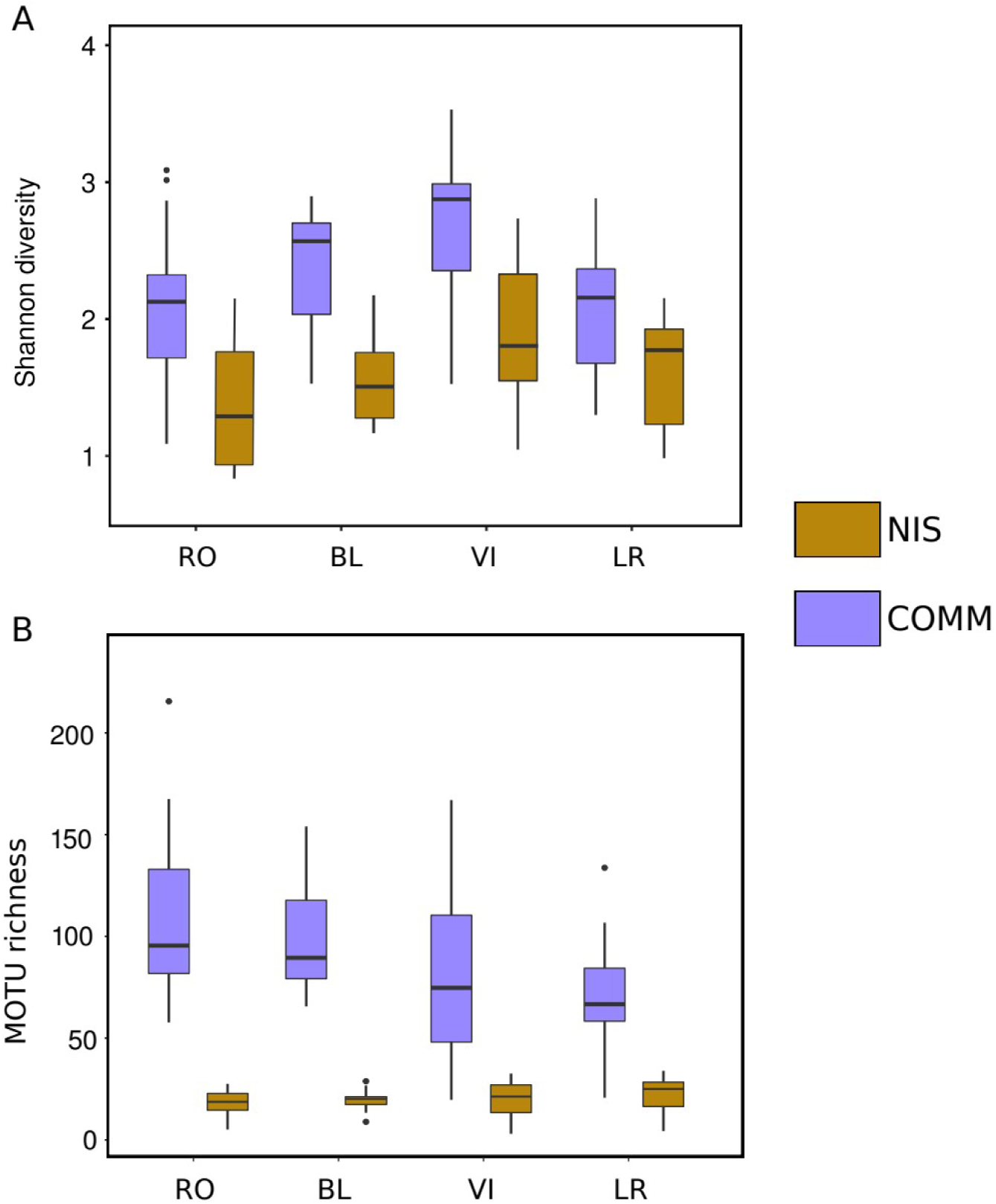
Box-plots of the values of Shannon diversity (A) and MOTU richness (B) of the COMM and NIS datasets. Horizontal lines are medians, boxes encompass the first and third quartiles, whiskers indicate 10th and 90th percentiles, and outliers are indicated as dot symbols. Site abbreviations are: RO - Roses, BL - Blanes, VI - Vilanova i La Geltrú, LR – La Ràpita.

The analyses of variance showed a significant effect of locality for both variables in both datasets (Table S3A). For the Shannon diversity index, post-hoc (Tukey) tests revealed significant differences in the COMM dataset for the comparisons VI-RO, and LR-VI. For the NIS dataset, the lowest diversity values across ports were found in the northernmost port (RO), while VI had the highest diversity. No comparison between ports was significant except VI with RO. For MOTU richness, RO had the highest values across ports in the COMM dataset, and the southern ports of VI and LR the lowest. The post-hoc tests revealed significant differences in the COMM dataset for the LR-RO comparison and showed no significant comparisons for the NIS dataset.

### 3.4. Distribution patterns and β-diversity

The relative read abundances of NIS were lowest in the northernmost port (RO) and highest in the southernmost port (LR), with intermediate values in BL and VI (Fig. 3). NIS always constituted more than 34% of the relative read abundance in each port, reaching over 70% in LR (Fig. 3A). However, in terms of the number of MOTUs, the proportion of NIS was minor compared to the total MOTUs of the entire community in all cases (Fig. 3B). In the locality with the highest proportion of NIS MOTUs (LR), they reached only 10% of the total. There was again an increasing trend from north to south, with the lowest proportion of NIS MOTUs in Blanes (7%). Figure 4 depicts the abundance of each NIS (MOTUs belonging to the same nominal species pooled) across localities, again reflecting their higher abundance in the southern ports.

**Fig. 3.**
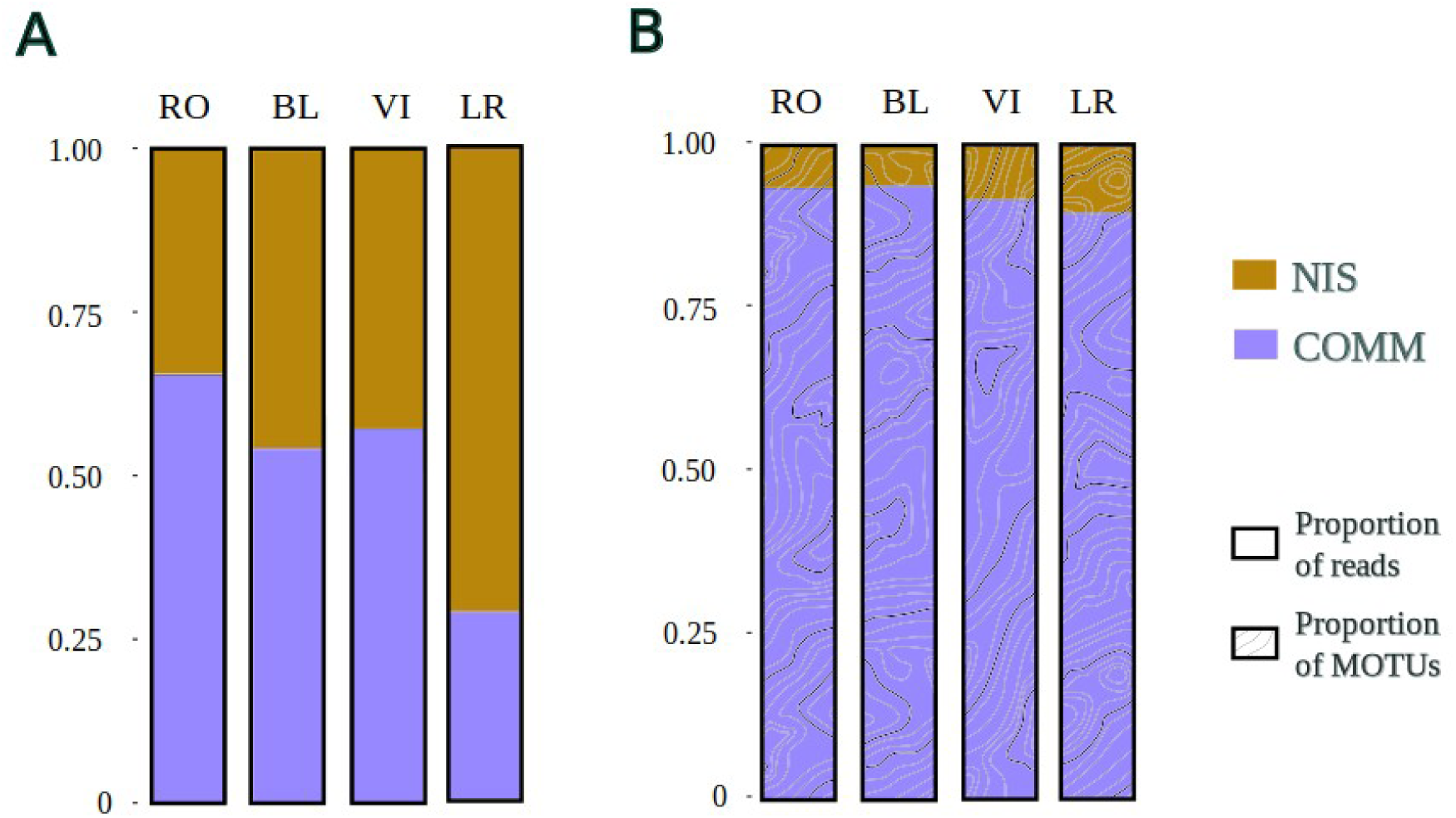
Barplots of the relative proportion of reads (A) and operational taxonomic units (MOTUs) (B) for the COMM and NIS datasets.

**Fig. 4.**
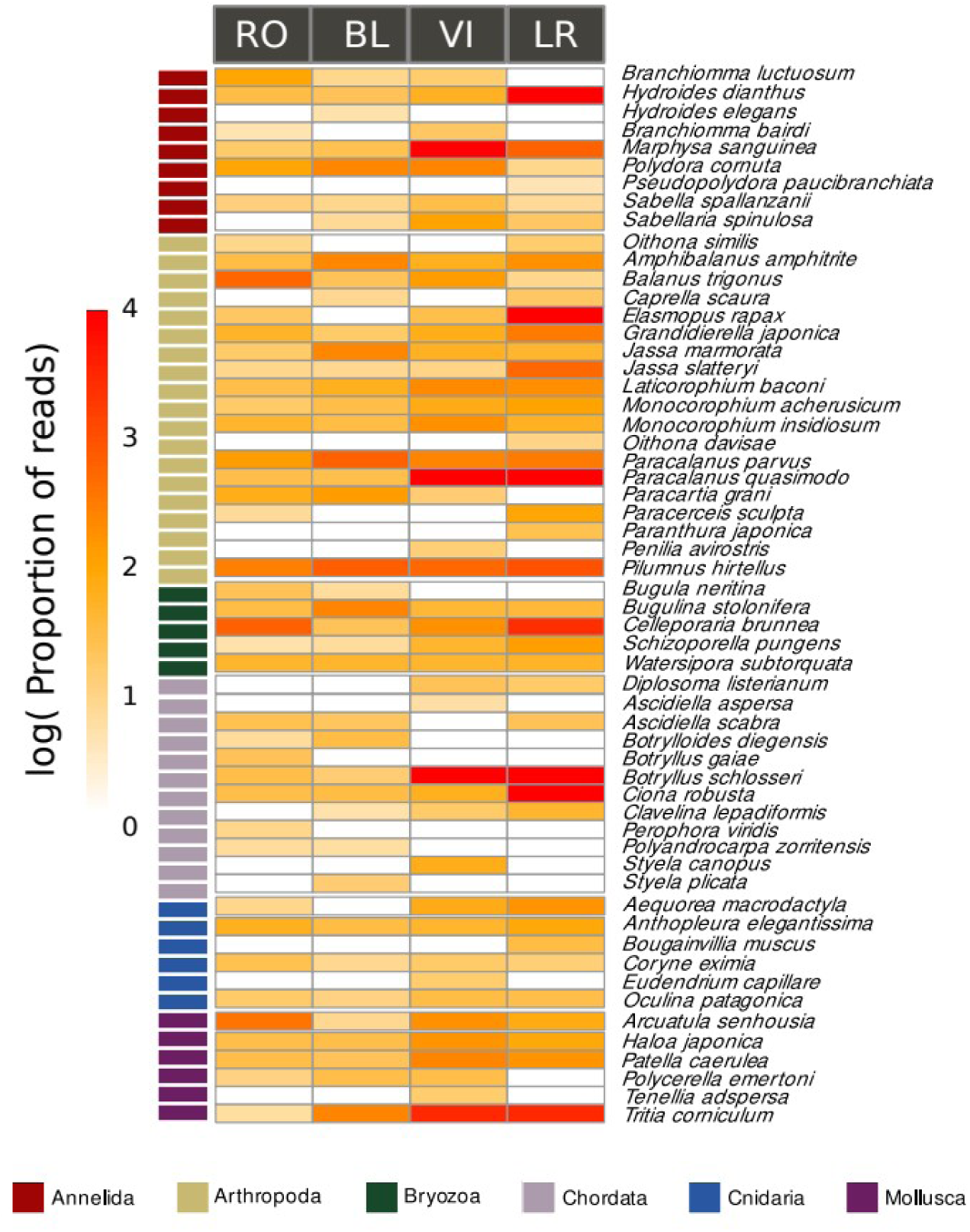
Heatmap showing relative read abundances (log-transformed) of the identified NIS in each area. Note that we have pooled molecular operational taxonomic units (MOTUs) assigned to the same nominal species. The phyla of the NIS are coded by colour.

Upset plots with the number of shared MOTUs (Fig. S6), showed markedly different patterns between COMM and NIS datasets. The majority of MOTUs of the former (66.7%) were unique to each locality (Fig. S6A). In contrast, the majority of NIS MOTUs (62.2%) were shared between at least two localities, and ca. one third were present in all four zones (Fig. S6B). The abundance of shared MOTUs between the different pairs of localities was also assessed with the match index (Fig. 5). The NIS MOTUs had a mean index of 0.828 among locality pairs, while the COMM MOTUs had a lower value (0.537), indicating less MOTU sharing, and the difference was significant (paired-sample *t*-test, p>0.001).

**Fig. 5.**
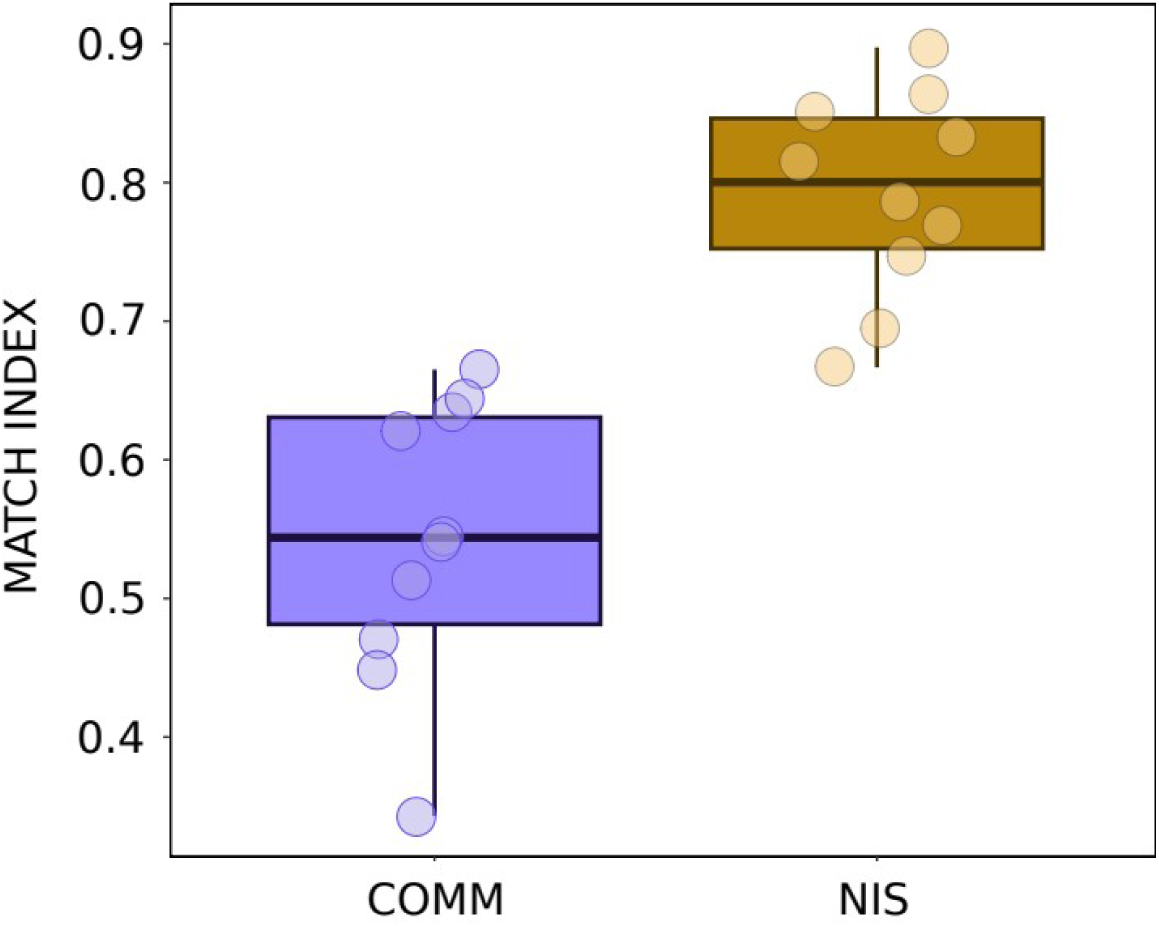
Box-plots of match index values for COMM and NIS MOTUs. Horizontal lines are medians, boxes encompass the first and third quartiles, whiskers indicate 10th and 90th percentiles.

Regarding the nMDS for the COMM datasets (Fig. 6A), the ordination along the first axis separated the four ports into two groups, the northern ports (RO and BL) and the southern ones (VI and LR). For the NIS dataset (Fig. 6B), the overall pattern was similar, but featuring less distinct clusters and more overlap. The mean BC dissimilarity values between pairs of samples was significantly higher for COMM than for NIS (mean of 0.776 and 0.688, respectively, paired-sample *t-*test, p<0.001).

**Fig. 6.**
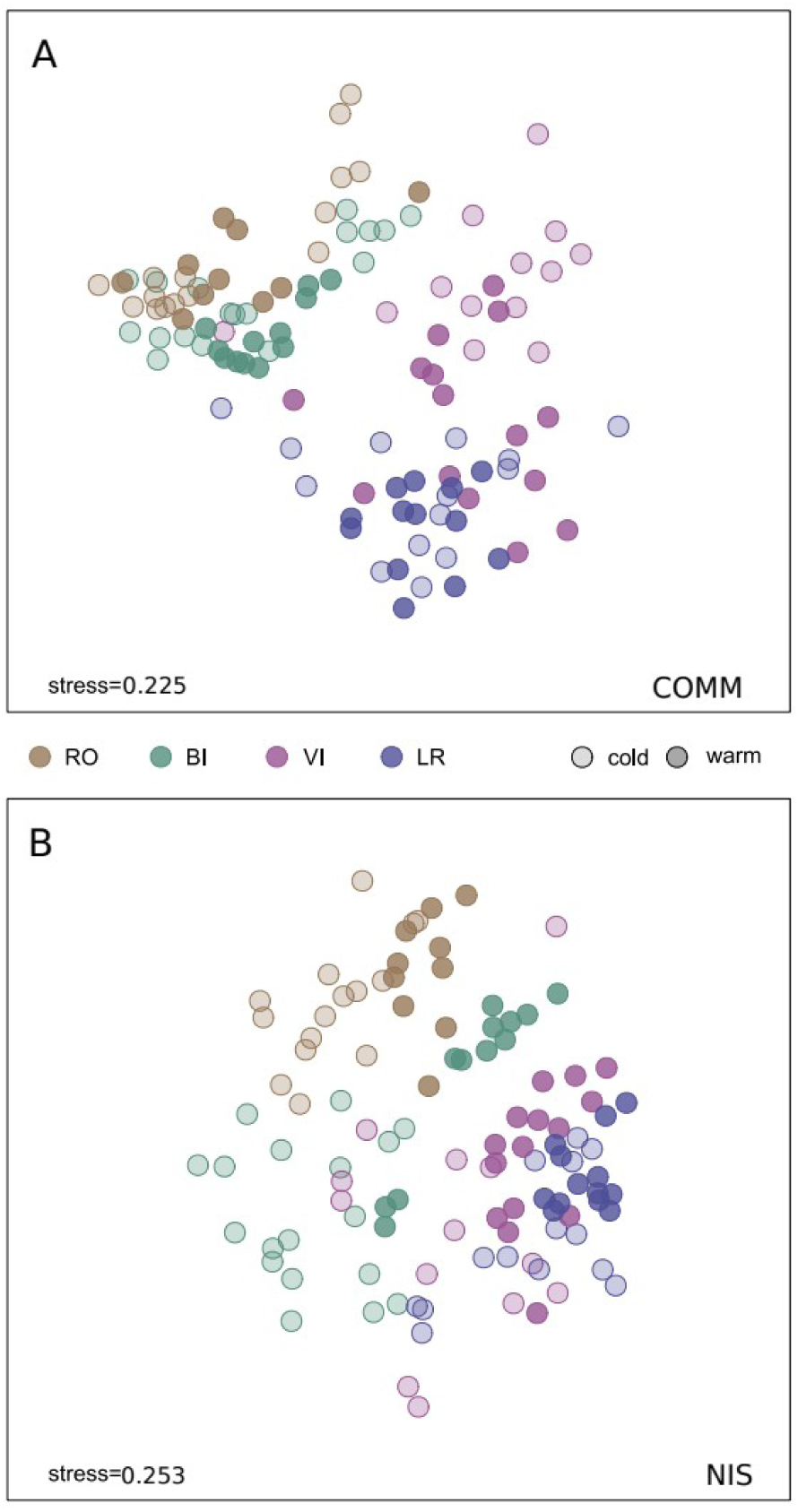
Non-metric MDS configurations for COMM (A) and NIS (B) datasets. Localities are indicated in different colours, and shading differentiates cold from warm months (applying a threshold of 20°C, solid colours corresponding to warm periods). Stresses of the final configurations are indicated.

The PERMANOVA tests indicated significant effects of site and month for both COMM and NIS MOTUs (Table S3B). The permdisp tests revealed significant differences in multivariate dispersion for the site factor in both datasets, but not for the month factor. Pairwise tests showed significant differences in both cases for all pairwise comparisons between sites. For the month factor, the COMM dataset had more significant pairwise comparisons than the NIS dataset. July was the month most differentiated from all others in both datasets. Consecutive sampling times were not significantly differentiated, except for June-July in both datasets and July-August in the NIS dataset.

Temperature of the sampled localities (Fig. S2) showed clear cut seasonal patterns, with lower summer temperatures in the northern ports (RO, BL), but lower winter temperatures in the South (CP, VI). Accordingly, some separation of cold and warm months, albeit with overlap, is also apparent in the MDS configurations for each site, both for COMM and NIS datasets (Fig. 6). We also plotted the mean Bray-Curtis dissimilarities values between consecutive sampling times separately for the COMM and NIS datasets (Fig. S7). For NIS these values peaked from July to February, indicating a higher species turnover, and were lowest in spring-beginning summer, suggesting more stability. The COMM dataset had overall higher BC distances and a similar temporal pattern.

### 3.5. Intra-MOTU genetic diversity and metaphylogeography

Fig. 7A shows the distribution of the number of ESVs per MOTU (log-transformed) in the NIS and NAT datasets. The 69.8% of the NIS MOTUs had some genetic variation, i.e., more than one ESV. This percent was lower (54.2%) for the NAT dataset. The median number of ESVs per MOTU was 7.39 for the NIS and 2.72 for the NAT MOTUs, and the difference was significant (Mann-Whitney test, p<0.001). The number of ESVs per MOTU was also significantly higher in the NIS dataset when the data were separated by locality (Fig. 7B, Mann-Whitney tests, all p<0.003). A higher ESV abundance in NIS MOTUs was detected also when the datasets were divided by phylum (Fig. 7C).

**Fig. 7.**
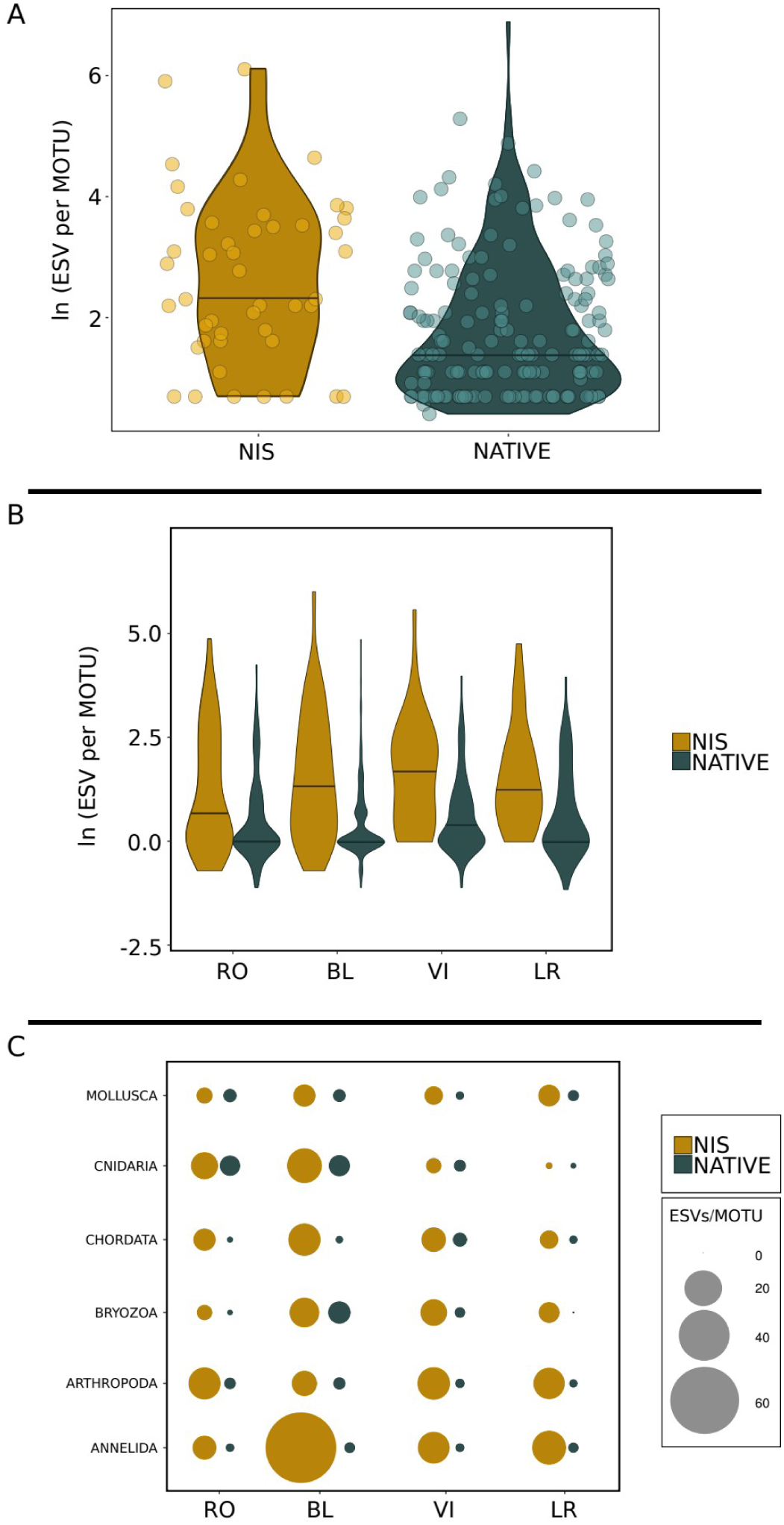
(A) Violin plots of the mean proportion of ESV per MOTU for NIS and native species. (B) violin plots separated by locality. (C) bubble plots as per phyla at each locality. Note natural logarithms in y axes of (A) and (B).

We obtained mean genetic differentiation (D) values between pairs of localities by averaging the D values of all MOTUs shared by each pair. These values (Fig. 8A) showed a lower differentiation in non-indigenous MOTUs (mean D=0.195) than in native MOTUs (mean D=0.314), and the difference was significant (paired-sample *t*-test, p=0.005). When the genetic distance measures for NAT and NIS were compared for each pair of localities (Fig. 8B), there was no significant correlation (*r*=0.19, p=0.6), indicating different patterns of genetic differentiation.

**Fig. 8.**
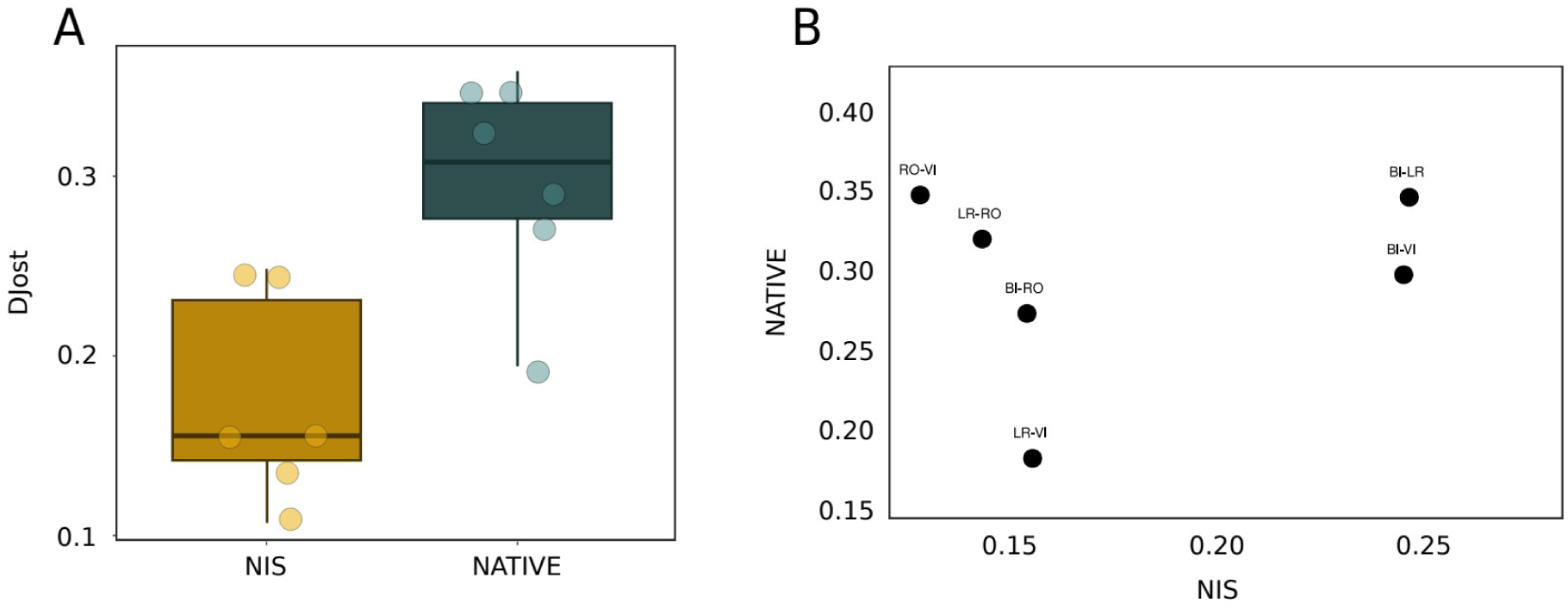
(A) Box-plots of the mean values of Jost’s D between localities for NIS and COMM datasets. Horizontal lines are medians, boxes encompass the first and third quartiles, whiskers indicate 10th and 90th percentiles. (B) Correlation plot for Djost for NIS and COMM datasets.

## 4. DISCUSSION

Our study showed that non-indigenous species exhibited higher population connectivity (considering both beta-diversity and genetic differentiation) and intraspecific genetic diversity compared to native species. These results support our initial expectation that maritime activities enhance NIS dispersal. The levels of genetic diversity detected in NIS suggests the presence of high propagule pressure in ports contributes to their invasion success, fostering higher genetic variation within these artificial communities. In contrast, native species display more restricted population connectivity and lower intraspecific genetic diversity, reflecting their isolation and vulnerability to impacts such as species invasions. These findings confirm that ports facilitate NIS’ range expansions, while the dispersal of native species among ports remains limited. By adopting a community-level approach and comparing native and NIS, our study provides valuable insights into population connectivity dynamics in modified ecosystems, an area that remains underexplored in the scientific literature, and provides a valuable baseline for future comparative studies.

DNA metabarcoding data from different substrates (water, settlement plates, sediment) and genetic markers offer different windows on port biodiversity, and combining them is recommended for detailed studies (Koziol et al., 2019; Lavrador et al., 2024; Rey et al., 2020). For broad-scale spatio-temporal comparisons of metazoan diversity, a cost-effective and practical approach is to use a standardised method such as the POMPOM collectors (Zarcero et al., 2024). Rarefaction curves indicated that our sequencing depth was sufficient to capture the diversity in each sample, although accumulation curves suggest that additional replicates could have revealed rare MOTUs, something common in metabarcoding studies (McIlroy et al., 2024; Turon et al., 2023; Wangensteen et al., 2018). Thus, we detected a substantial proportion of the biodiversity present in the studied ports that can be used to confidently assess patterns of population connectivity and intraspecific genetic diversity.

Interestingly, while NIS represented less than 4% of the total MOTUs, they had a disproportionate abundance in terms of number of reads, comprising between 34 and 70% of the reads in the port samples. In another metabarcoding study of harbour communities, Lavrador et al. 2024 detected a similar percentage in species richness (3.6%) of NIS using several substrate types. Arthropods, cnidarians and chordates (ascidians) dominated port communities in our samples, and a gradient was detected as the abundance of cnidarians decreased and that of arthropods increased from north to south. No similar gradient was observed in the taxonomic composition of the NIS found in the ports, dominated by arthropods, annelids and chordates.

The southernmost ports (VI and LR) displayed an overall lower MOTU richness, and LR had the highest abundance and diversity of NIS. It is unclear which factor or combination of factors can drive these differences. Temperature may play a role, as readings indicated a warmer summer, but colder winter, in the southernmost site. In addition, our sampling points are interspersed with two of the largest commercial ports in Western Mediterranean, namely Barcelona (between BL and VI) and Tarragona (between VI and LR). These large harbours are home of intense international cruise and cargo trade operations and act as entry points for species and subsequently influence the composition of nearby smaller ports via local boating. However, the higher presence of NIS in LR is more likely explained by the proximity of this port to the lagoons of the Ebro Delta, one of the largest bivalve aquaculture regions of the Mediterranean Sea. Indeed, the Ebro Delta area is a NIS hotspot (Casso et al., 2018) and can thus influence the species composition of surrounding areas.

Our MDS configurations for overall community composition in ports showed a north to south gradient. Interestingly, all four ports clustered separately (with some overlap) in the final configuration, highlighting significant spatial structure at this relatively small scale. In line with this, the MDS analysis of the MOTUs identified as NIS indicated a similar overall trend, but with a significantly higher overlap and less distinctness of the communities in each locality studied. The finding of weaker spatial structure in the MOTUs assigned to NIS, compared to the overall community, was consistent with the greater MOTU sharing observed in the NIS dataset. In fact, 36.6% of the NIS MOTUs were detected in all study areas, whereas 66.7% of the MOTUs in the COMM dataset were restricted to only one of the four localities. This contrasts with the 6.4% of COMM MOTUs that were found across all four localities. Likewise, the match ratio among localities was significantly higher for the NIS dataset. Boat traffic likely fuels this differential connectivity of species well adapted to travel as hitchhikers on hulls (Ulman et al. 2019a). For most other species, coastal drift or other natural dispersal processes are the only way to ensure some degree of connectivity (Cowen et al 2007).

We identified some interesting temporal patterns in our samples, such as the quantitative replacement of some groups over the studied months, both for NIS and for the wider community. Common NIS occurring along the Catalan coast are known to show highly seasonal patterns (e.g. Rius et al., 2009), while other NIS are present and reproductive all year around (e.g. Pineda et al., 2013). We also detected differences in beta-diversity analyses such as clustering in cold or warm periods of the samples.. Previous metabarcoding studies have also detected seasonal changes in port communities (Rey et al., 2020, Lavrador et al., 2024).

Our use of a denoising method customised for coding sequences to generate an ESV dataset allowed us to uncover intra-MOTU patterns. In this context, we used ESVs as a proxy for haplotypes in the same way that MOTUs are a proxy for species. To ensure a more targeted comparison, we selected a subset of MOTUs from the COMM dataset that could be confidently assigned to native species. We focused our analyses on these native MOTUs rather than the rest of the community, as MOTUs not identified at species level may include cryptogenic or unrecognised NIS, potentially biasing intraspecific diversity estimates. We then examined their composition in terms of ESVs within MOTUs as compared with the NIS dataset. This comparison allowed an assessment of the scope for adaptation of these groups. A first analysis showed that the genetic richness was significantly higher in NIS compared to NAT. In addition, this holds true at a phylum-by-phylum basis, showing that this effect is not restricted to a particular group behaving differently. This pattern was somewhat unexpected, as NIS are supposed to experience bottlenecks when they colonise new areas. The so-called genetic invasion paradox (Roman and Darling, 2007) has long been a matter of debate, mainly when NIS thrive when their genetic variability is drastically reduced. This reduction may simply be a wrong assumption, as recurrent introduction of propagules from different regions can in fact lead to higher genetic diversity in introduced than in natural populations of these species, thus solving the paradox (Estoup et al., 2016; Lacoursière-Roussel et al., 2016; Roman and Darling, 2007). In contrast, native species may rely on a more limited source of new propagules in ports (Allendorf and Lundquist, 2003) and may become comparatively haplotype-poor. Finding suitable environments combined with high genetic diversity may be key to their successful settlement, survival, and dispersal strategies (Roman and Darling, 2007), which is key for eventually becoming a biological invasion problem (Miralles et al., 2016). Genomic features, such as introgressions (Touchard et al., 2024), inversions (Galià-Camps et al., 2024), or the microbiome (Casso et al., 2020; Galià-Camps et al., 2023) have been proposed as possible explanations for the fast adaptation of NIS to new environments, and a high genetic variability can also contribute to their ability to do so.

Our finding of high genetic connectivity observed among non-indigenous species in Mediterranean ports showed evidence of intense gene flow across multiple locations, contrasting with the more marked spatial structure observed in native species. This high level of connectivity likely promotes genetic homogenisation and helps maintaining elevated intraspecific diversity, which is known to enhance the adaptive capacity and invasive success of NIS. As seen in previous studies, our results confirm the role of ports as networks of propagation of NIS (López-Legentil et al 2015) and the existence of a portuarization syndrome (Touchard et al., 2022, Lilli et al., 2025), whereby ports act as replicate natural laboratories favouring the evolution of well-adapted, homogenised NIS biota. This phenomenon not only fuels biological invasions but also poses a critical challenge for the management and conservation of coastal ecosystems, as high genetic diversity and high adaptive capabilities in NIS may increase their resilience to environmental changes and their potential to spread and displace of native communities.

## 5. CONCLUSION

Our results revealed a high population connectivity of NIS across a network of medium-sized ports in the north-western Mediterranean, highlighting their pivotal role in structuring and connecting port ecosystems. At the community level, NIS displayed distinct composition and dynamics, characterised by pronounced homogenisation among ports compared to the rest of the assemblage. This pattern, reflected in lower beta-diversity and higher MOTU sharing, suggests that NIS are more efficiently dispersed, most likely facilitated by recreational and fishing vessel movements linking these harbours. When comparing NIS with native species, similarly striking genetic differences emerged. NIS exhibited significantly lower population differentiation and greater haplotype richness, indicating more intense gene flow and recurrent lineage exchange among ports, promoted by continued propagule introductions. In contrast, native species displayed more fragmented population structure and lower intraspecific genetic diversity, reflecting higher levels of isolation, and hence vulnerability, in highly anthropogenic coastal environments. These findings fully support our initial hypotheses: NIS are more homogeneous among ports, exhibiting stronger genetic connectivity and lower differentiation, and maintain higher intraspecific genetic diversity than native species. Moreover, the high genetic variability observed in NIS challenges the classical “genetic invasion paradox”, which predicts diversity loss through population bottlenecks during colonisation. Our results instead suggest that recurrent propagule introductions from multiple source regions can enhance genetic diversity in invasive populations, thereby promoting their adaptive potential and ultimately, their invasion success.In addition, this study highlights the value of integrating inter- and intra-MOTU analyses to understand invasion processes, from community assembly to genetic differentiation. Finally, our results suggest the need for both well-curated genetic reference databases and continued monitoring in medium-sized harbours, which may act as critical nodes for the secondary spread of NIS.

### CRediT AUTHORSHIP CONTRIBUTION STATEMENT

**Jesús Zarcero**: Writing – original draft, Writing – review and editing, Data curation, Formal analysis, Investigation, Methodology, Software, Visualization. **Adrià Antich**: Writing – review and editing, Conceptualization, Data curation, Software. **Margarita Fernández**: Writing – review and editing, Methodology. **Cruz Palacín**: Writing – review and editing, Conceptualization, Methodology. **Owen Wangensteen**: Writing – review and editing, Project administration, Resources, Software, Supervision, Validation. **Marc Rius**: Writing – review and editing, Funding acquisition, Project administration, Resources, Supervision, Validation. **Xavier Turon**: Writing – original draft, Writing – review and editing, Conceptualization, Data curation, Formal analysis, Funding acquisition, Investigation, Methodology, Project administration, Resources, Software, Supervision, Visualization.

## ACKNOWLEDGMENTS

The Department of Territory of the Catalan Government (“Ports de la Generalitat”) granted the permissions to work in the harbours surveyed. We thank Paco Falcó who provided access to his mooring place in Roses to place the collectors. Thanks also to the skippers of IRTA for accompanying us during the samplings in La Ràpita.

---

**Fig. S1.**
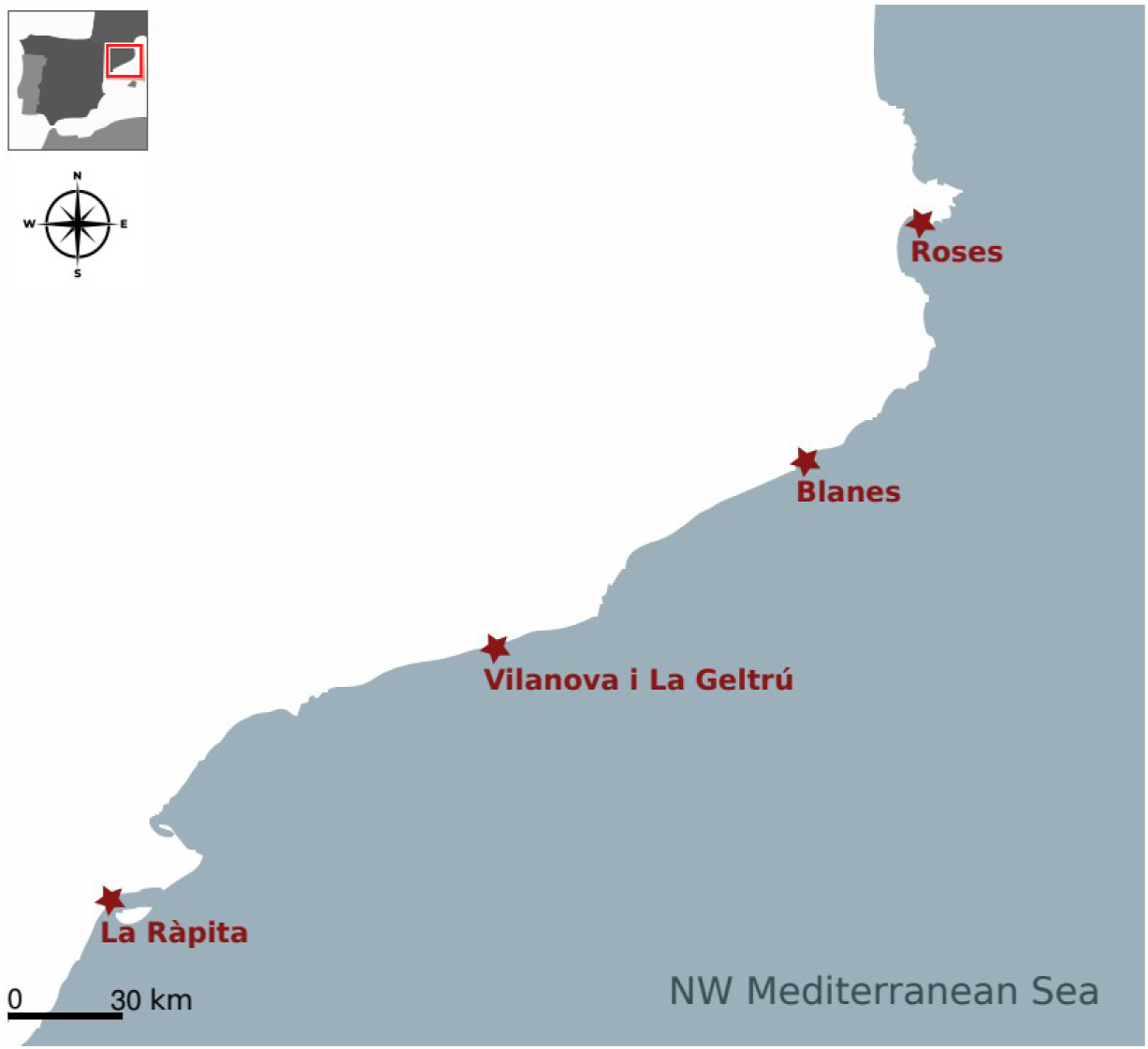
Map of the study area (Catalonia, Spain, NW Mediterranean) indicating each locality where samples were collected.

**Fig. S2.**
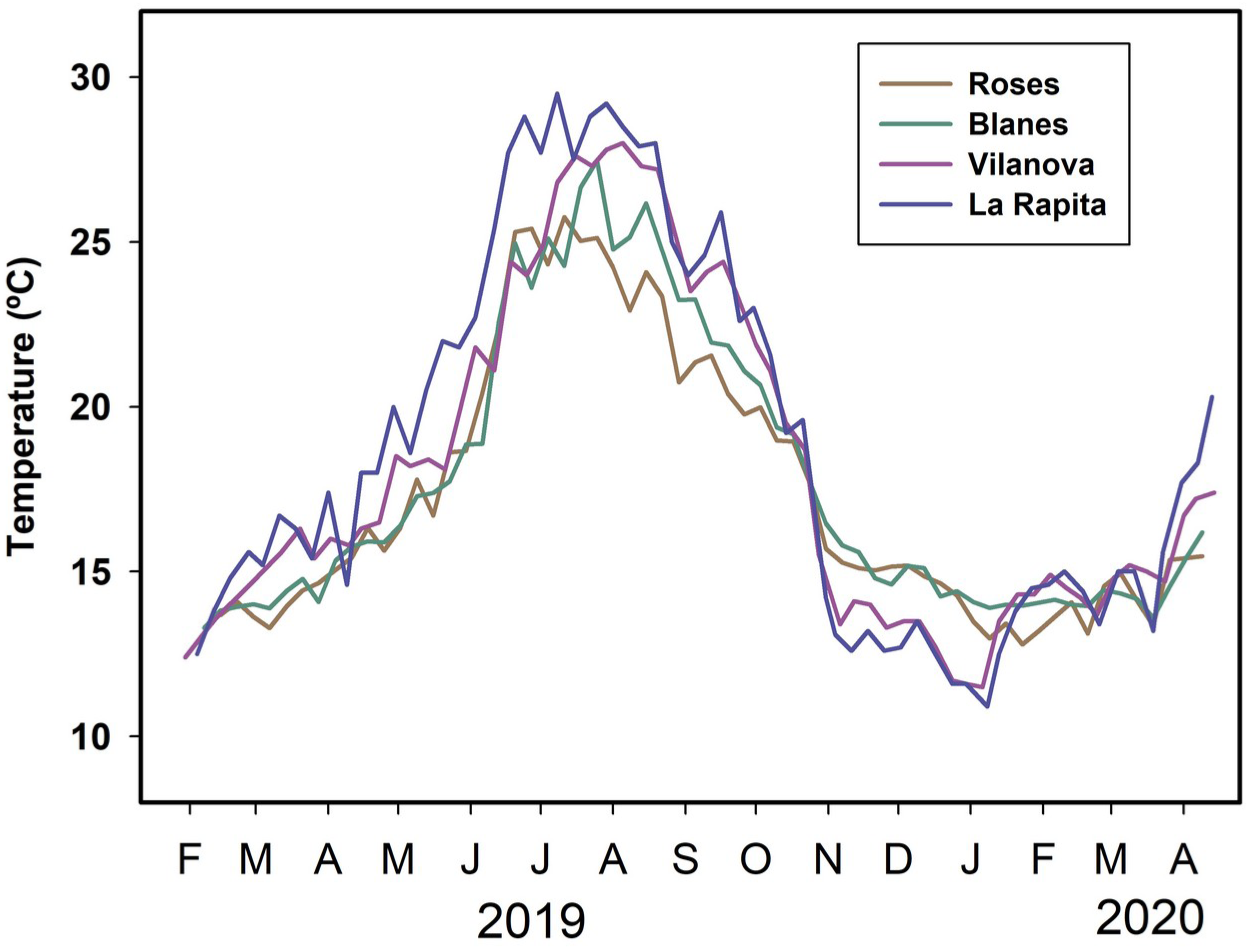
Graph of the water temperature values for each area during the study period (from February 2019 to April 2020).

**Fig. S3.**
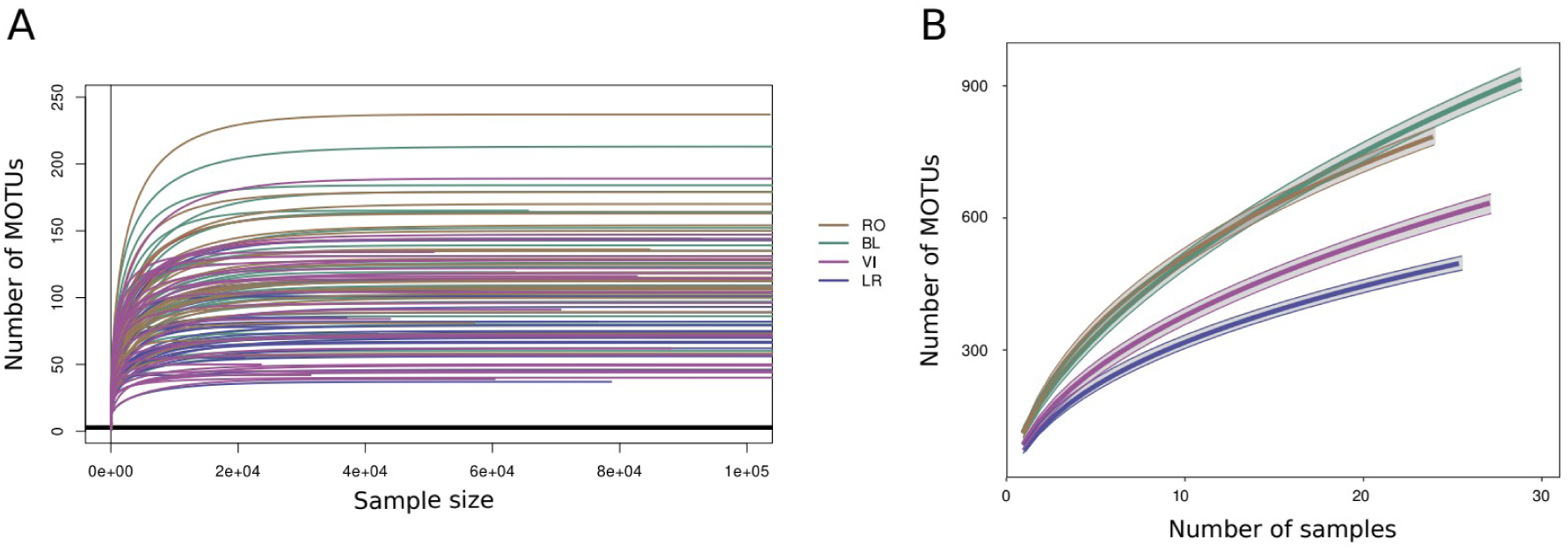
(A) Rarefaction curves indicating the number of MOTUs recovered at increasing read depth (up to 100,000 reads for clarity) in the samples (RO: Roses; BL: Blanes; VI: Vilanova i La Geltrú; LR: La Ràpita). (B) MOTU accumulation curves at increasing number of samples. Accumulation curves are displayed per area, corresponding to all metazoan MOTUs detected.

**Fig. S4.**
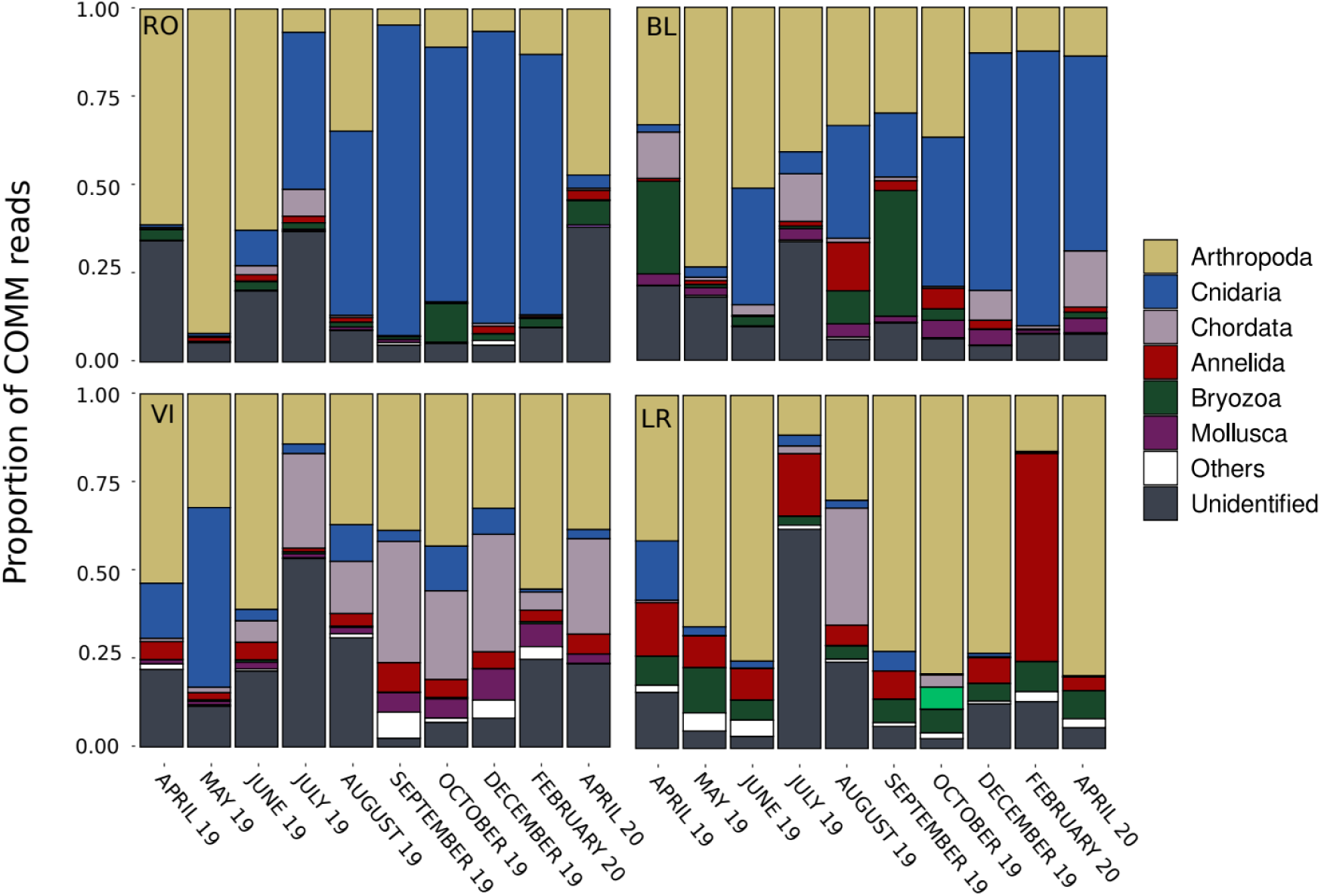
Barplots of the relative proportion of COMM reads for each locality (RO: Roses; BL: Blanes; VI: Vilanova i La Geltrú; LR: La Ràpita) of the different phyla considered at each sampling time.

**Fig. S5.**
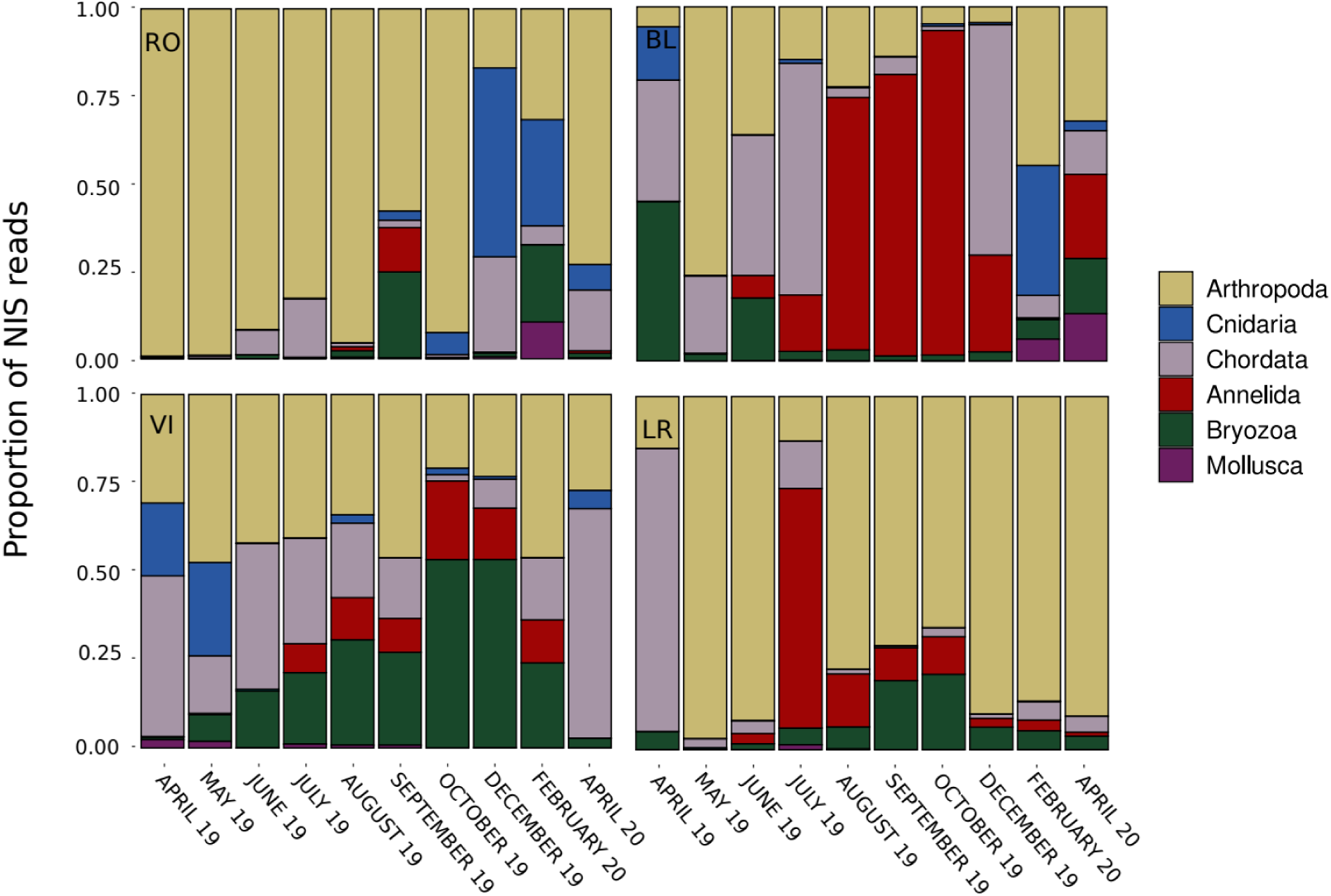
Barplots of the relative proportion of NIS reads for each locality (RO: Roses; BL: Blanes; VI: Vilanova i La Geltrú; LR: La Ràpita) of the different phyla considered at each sampling time.

**Fig. S6.**
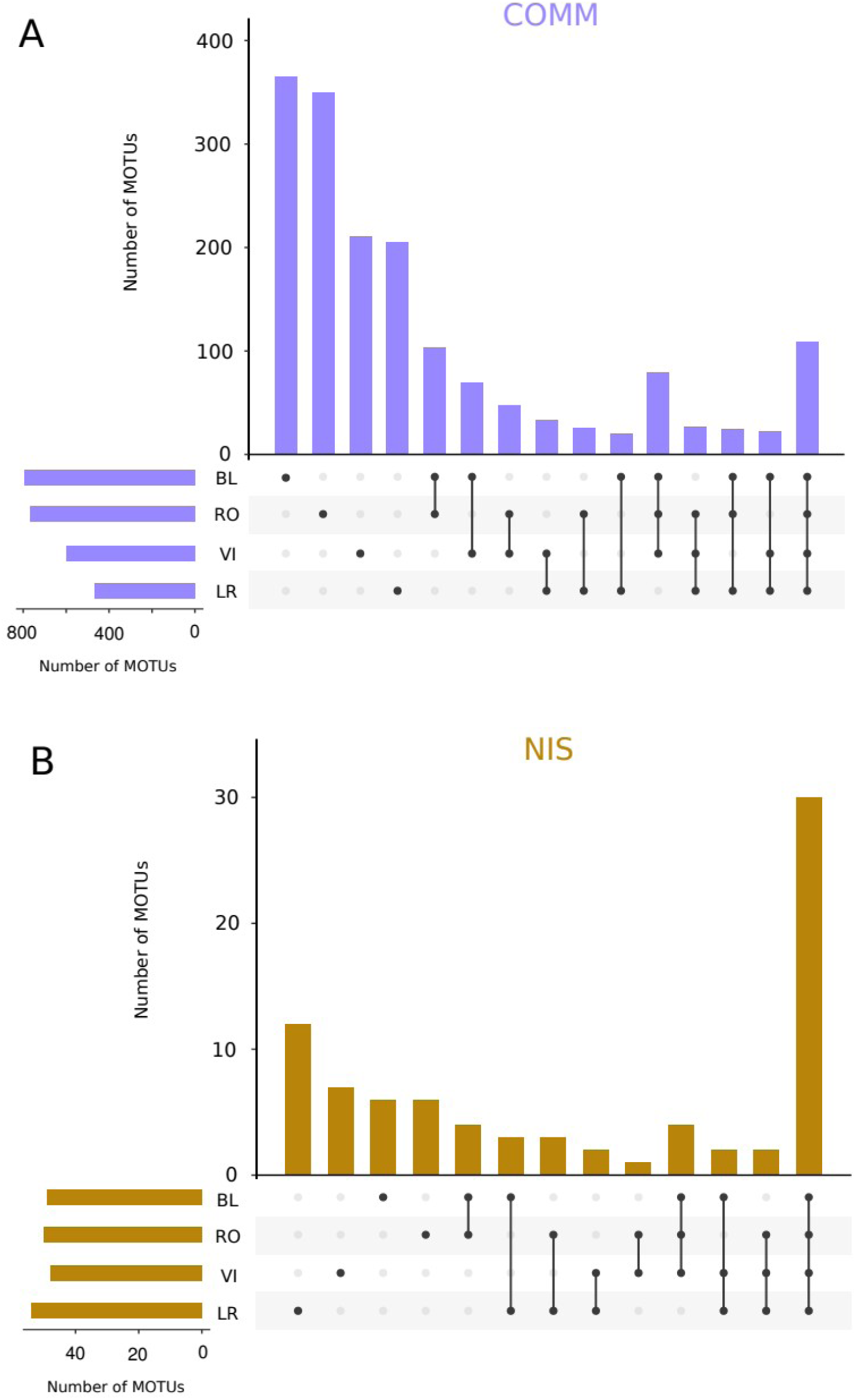
Upset-plots of shared MOTUs between localities for COMM (A), and NIS (B).

**Fig. S7.**
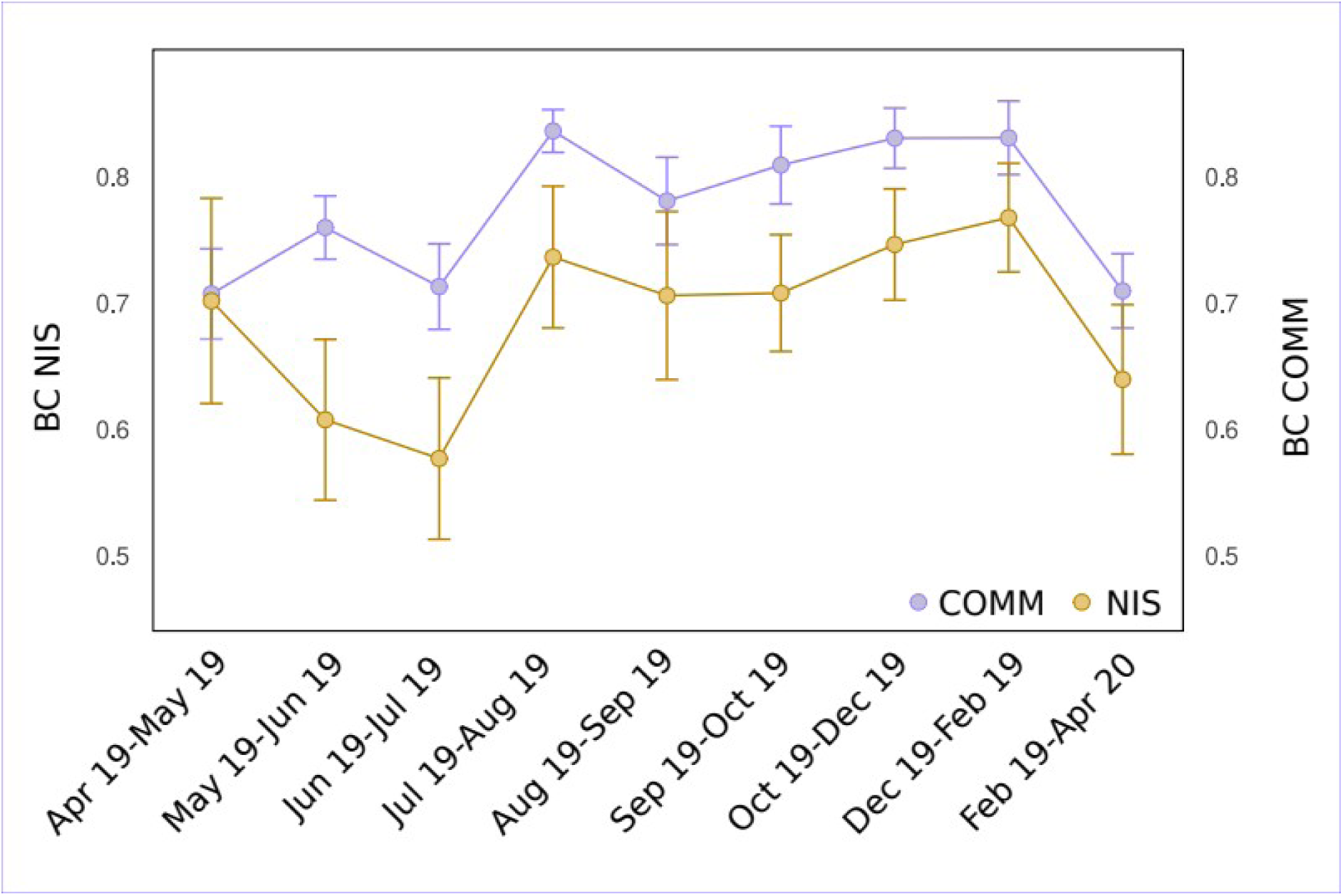
Mean values of Bray-Curtis distance for consecutive sampling times for COMM and NIS datasets. The supplementary tables are available at: https://github.com/jesuszarcero/RBVC_supplementary_data

**Table S1.** Table of the 1,774 metazoan MOTUs identified in the analyses. For each MOTU the following information is provided: id, pid, name, taxonomic rank at which it could be assigned, taxonomic assignments at the different levels, total abundance in number of reads, abundance in each sample (see “Sample metadata” sheet), representative sequence of the MOTU, and identification of the corresponding category: NIS and COMM (in the latter with indication of MOTUs labelled as native). For NIS species, the identification obtained with the custom NIS database is also presented.

**Table S2.** Table of the 11,193 metazoan ESVs used in the analyses. For each ESV the following information is provided: id, total abundance in number of reads, MOTU to which it belongs, abundance in each sample (see “Sample metadata” sheet), and sequence..

**Table S3.** (A) Table with ANOVA tests and Tukey tests for the Shannon diversity and MOTU richness variables. (B) Results of the PERMANOVA tests and pairwise comparisons on the Bray-Curtis dissimilarity matrix.

